# Needle in a haystack? A comparison of eDNA metabarcoding and targeted qPCR for detection of the great crested newt (*Triturus cristatus*)

**DOI:** 10.1101/215897

**Authors:** Lynsey R. Harper, Lori Lawson Handley, Christoph Hahn, Neil Boonham, Helen C. Rees, Kevin C. Gough, Erin Lewis, Ian P. Adams, Peter Brotherton, Susanna Phillips, Bernd Hänfling

## Abstract

Environmental DNA (eDNA) analysis is a rapid, cost-effective, non-invasive biodiversity monitoring tool which utilises DNA left behind in the environment by organisms for species detection. The method is used as a species specific survey tool for rare or invasive species across a broad range of ecosystems. Recently, eDNA and ‘metabarcoding’ have been combined to describe whole communities rather than focusing on single target species. However, whether metabarcoding is as sensitive as targeted approaches for rare species detection remains to be evaluated. The great crested newt *Triturus cristatus* is a flagship pond species of international conservation concern and the first UK species to be routinely monitored using eDNA. We evaluate whether eDNA metabarcoding has comparable sensitivity to targeted real-time quantitative PCR (qPCR) for *T. cristatus* detection. Extracted eDNA samples (N = 532) were screened for *T. cristatus* by qPCR and analysed for all vertebrate species using High-Throughput Sequencing technology. With qPCR and a detection threshold of 1/12 positive qPCR replicates, newts were detected in 50% of ponds. Detection decreased to 32% when the threshold was increased to 4/12 positive qPCR replicates. With metabarcoding, newts were detected in 34% of ponds without a detection threshold, and in 28% of ponds when a threshold (0.028%) was applied. Therefore, qPCR provided greater detection than metabarcoding but metabarcoding detection with no threshold was equivalent to qPCR with a stringent detection threshold. The proportion of *T. cristatus* sequences in each sample was positively associated with the number of positive qPCR replicates (qPCR score) suggesting eDNA metabarcoding may be indicative of eDNA concentration. eDNA metabarcoding holds enormous potential for holistic biodiversity assessment and routine freshwater monitoring. We advocate this community approach to freshwater monitoring to guide management and conservation, whereby entire communities can be initially surveyed to best inform use of funding and time for species-specific surveys.

## Introduction

Species monitoring has rapidly evolved with the advent of environmental DNA (eDNA) analysis (Lawson Handley 2015). eDNA analysis allows highly sensitive detection of rare and invasive species and is increasingly being used for surveys of aquatic species (Thomsen *et al.* 2012; Biggs *et al.* 2015; Davy, Kidd & Wilson 2015; Smart *et al.* 2016; Evans *et al.* 2017b). This non-invasive approach uses intra- and extracellular DNA (mucous, skin cells, faeces/urine, gametes, hair, deceased remains) released into the environment by organisms to survey for species and assess their distribution (Rees *et al.* 2014b; Lawson Handley 2015; Goldberg *et al.* 2016). Typically for eDNA analysis, DNA is extracted from environmental samples (water, soil, air) and analysed using a targeted or passive approach. The targeted approach uses species-specific primers with conventional PCR (PCR), real-time quantitative PCR (qPCR), or droplet digital PCR (ddPCR), to determine presence-absence and estimate abundance of single species (Shaw, Weyrich & Cooper 2016b; Goldberg *et al.* 2016). Conversely, the passive approach uses conserved primers (i.e. primers with binding sites that are shared across multiple taxa, and flank a region of highly variable DNA sequence that enables discrimination between these taxa) and PCR to sequence whole communities with High Throughput Sequencing (HTS), termed eDNA metabarcoding (Taberlet *et al.* 2012; Shaw *et al.* 2016b; Valentini *et al.* 2016; Deiner *et al.* 2017). Passive eDNA monitoring is particularly attractive to ecologists for biodiversity assessment as a means to detect entire species assemblages alongside rare or invasive species (Lacoursière-Roussel *et al.* 2016; Blackman *et al.* 2017). However, this gain in community understanding may come at the cost of accuracy and sensitivity. Direct comparisons of these two approaches are essential to determine whether they have comparable power and yield similar results.

Although in its relative infancy, eDNA metabarcoding has proven effective for community biodiversity assessment across a range of taxa in varying environments, particularly freshwater herpetofauna and fish (Valentini *et al.* 2016; Hänfling *et al.* 2016; Shaw *et al.* 2016a; Civade *et al.* 2016; Lacoursière-Roussel *et al.* 2016; Lopes *et al.* 2016; Evans *et al.* 2017a; Bálint *et al.* 2017). However, eDNA metabarcoding is confounded by potential amplification bias during PCR, preventing capture of all species present in a given area (Kelly *et al.* 2014). Species’ DNA in community samples is also in competition to bind to metabarcoding primers during PCR, where more common templates are more likely to be amplified. High abundance species may thus prevent detection of low abundance species, whether by fewer individuals or less DNA shed, resulting in ‘species masking’ (Kelly *et al.* 2014; Brandon-Mong *et al.* 2015; Evans *et al.* 2016). eDNA metabarcoding may therefore be less capable of identifying eDNA of rare species within a community than species-specific qPCR (Evans *et al.* 2016).

The sensitivity of eDNA metabarcoding has been evaluated against conventional biodiversity monitoring methods in freshwater ecosystems (Valentini *et al.* 2016; Hänfling *et al.* 2016; Shaw *et al.* 2016a; Civade *et al.* 2016; Lopes *et al.* 2016; Evans *et al.* 2017a; Bálint *et al.* 2017), yet specific investigations comparing the sensitivity of eDNA metabarcoding and targeted qPCR are sparse. Similarly, comparisons of qPCR and conventional survey for species monitoring have included cost projections (Biggs *et al.* 2014; Davy *et al.* 2015; Smart *et al.* 2016; Evans *et al.* 2017b) but cost has not been thoroughly assessed in qPCR and eDNA metabarcoding comparisons (Lacoursière-Roussel *et al.* 2016; Schneider *et al.* 2016). Schnieder *et al.* (2016) achieved improved detection of Invasive Mosquito Species (IMS) with qPCR and eDNA metabarcoding as opposed to conventional sampling. Although qPCR provided higher detection probability for two species, metabarcoding achieved comparable results for a third species, and allowed simultaneous detection of IMS and other taxa in a single sequencing run without development of multiple species-specific markers. In another study, eDNA metabarcoding failed to detect wood turtle *Glyptemys insculpta* (LeConte, 1830) in four rivers where qPCR and conventional visual survey detected the species (Lacoursière-Roussel *et al.* 2016). Amplification of longer fragments during metabarcoding versus qPCR could account for difference in sensitivity of the two methods, with the shorter qPCR assay being more capable of detecting heavily degraded DNA (Lacoursière-Roussel *et al.* 2016). Further research is clearly needed to determine whether these two approaches are comparable.

Great crested newt *Triturus cristatus* (Laurenti, 1768) (Fig. 1) is a model organism for eDNA-based monitoring. *T. cristatus* secrete mucous, breed in water, and produce aquatic eggs and larvae all sources of DNA deposition in ponds. The species is rare in parts of the UK and Europe, and as such all life stages are protected by UK and European legislation (Rees *et al.* 2014a; Buxton *et al.* 2017). eDNA analysis using targeted qPCR has been repeatedly verified against conventional surveying (bottle trapping, torchlight counts, larval netting, egg searches) for *T. cristatus* and found to achieve comparable or improved species detection (Thomsen *et al.* 2012; Rees *et al.* 2014a; Biggs *et al.* 2015). eDNA sampling can be undertaken with relative ease, is cost-efficient (Biggs *et al.* 2014) and can be implemented in large-scale citizen science monitoring programmes without loss of species detection (Biggs *et al.* 2015). *T. cristatus* is the first species to be routinely monitored using eDNA in the UK (Natural England 2015) and targeted eDNA assays are now offered as a commercial service by ecological consultancies. The targeted eDNA assay is highly effective for *T. cristatus* detection; however, should metabarcoding have comparable sensitivity, this approach would allow detection of *T. cristatus* alongside pond communities and potentially enable more cost-effective monitoring of entire ecosystems and ecological hypothesis testing.

**Figure 1.**
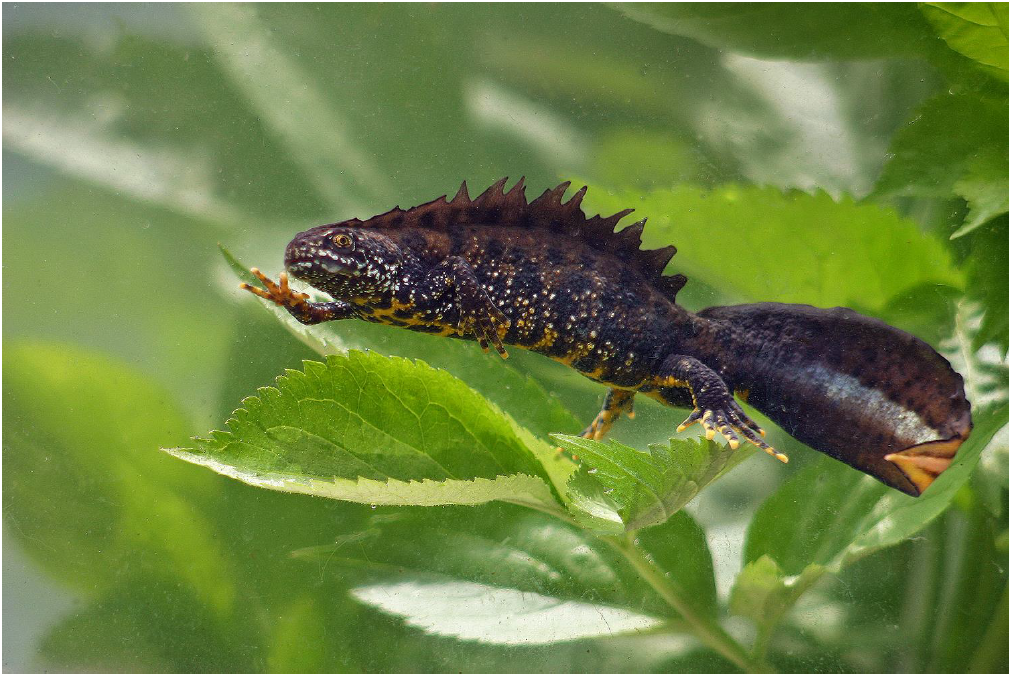
Adult male great crested newt (*Triturus cristatus*). Photo by Brett Lewis (Lewis Ecology, Brett Lewis Photography).

Here, we perform a large-scale comparison (N = 532 ponds) of eDNA metabarcoding and targeted qPCR for *T. cristatus* detection to compare method sensitivity. A single primer pair that is vertebrate specific for mitochondrial DNA (mtDNA) and requires no *a priori* knowledge of species composition was employed for eDNA metabarcoding. The metabarcoding results were then compared to results obtained using the standard *T. cristatus* qPCR assay (Biggs *et al.* 2015). Our hypotheses are: (1) eDNA metabarcoding will give equivalent results to qPCR for *T. cristatus* detection, (2) eDNA metabarcoding sequence read count for *T. cristatus* will increase as qPCR score (the number of positive qPCR replicates) increases, indicative of eDNA concentration, and (3) metabarcoding primers will amplify DNA from all taxa equally well and no bias towards amplification of *T. cristatus* will occur (bias would be indicated by a positive association between the proportion of *T. cristatus* sequence reads and PCR product concentration). We also examined cost and investigator effort required by each approach to determine whether a trade-off between cost, time and amount of data generated exists.

## Materials and methods

### SAMPLING

Samples from 532 ponds distributed across three UK counties (Cheshire, Kent and Lincolnshire) were analysed for this project. Of these, 508 ponds (ranging from 9 to 9375 m^2^) were sampled as part of *T. cristatus* surveys through Natural England’s Great Crested Newt Evidence Enhancement Programme. *T. cristatus* egg searches were performed once during the daytime at 506/508 ponds. Any other life stages seen were also recorded. A further 24 ponds were sampled for eDNA by ecological consultants for private contracts but egg searches were not undertaken. All water samples were collected using methodology outlined by Biggs *et al.* (2015). Water samples were then sent to Fera (Natural England) and ADAS (private contracts), where one eDNA sample per pond was produced and analysed according to laboratory protocols established by Biggs *et al.* (2015). Details of sampling methodology and laboratory protocols are provided in Supporting Information: Appendix 1.

### TARGETED qPCR FOR *T. CRISTATUS*

Targeted qPCR was conducted as part of the *T. cristatus* monitoring programmes mentioned above in Fera and ADAS laboratories during 2015. Both laboratories used a standardised protocol, which tests for PCR inhibitors and sample degradation prior to testing for *T. cristatus* (Biggs *et al.* 2015).Extracted DNA was amplified by TaqMan probe qPCR using published primers and probe (Thomsen *et al.* 2012) to amplify an 81 bp fragment of the cytochrome b gene. For each sample, 12 qPCR replicates were performed and a sample recorded as positive for *T. cristatus* if one or more qPCR replicates were positive. Following qPCR, the eDNA samples were placed in storage at -80°C.

### METABARCODING OF VERTEBRATE COMMUNITIES

eDNA samples were stored at -20 °C until PCR amplification. Metabarcoding was performed using published vertebrate-specific primers (Riaz *et al.* 2011) which amplify a 73-110 bp fragment of 12S ribosomal RNA gene (rRNA). The assay was first validated *in silico* using ecoPCR software (Ficetola *et al.* 2010; Bellemain *et al.* 2010) against a custom, phylogenetically curated reference database for UK vertebrates. Full details of reference database construction are provided in Supporting Information: Appendix 2. The complete reference database compiled in GenBank format has been deposited in a dedicated GitHub repository for this study, permanently archived at: https://doi.org/10.5281/zenodo.1188709. Parameters set allowed a 50-250 bp fragment and maximum of three mismatches between the primer pair and each sequence in the reference database. Primers were then validated against tissue DNA extracted from UK amphibian species (Supporting Information: Appendix 2) having been previously validated *in vitro* for UK fish communities by Hänfling *et al.* (2016). After primer validation, a two-step PCR protocol was used to construct metabarcoding libraries from the eDNA samples. During the first PCR, the target region was amplified using metabarcoding primers, comprised of the aforementioned specific locus primer, random hexamers, sequencing primer and pre-adapter (Illumina 2011). DNA from the cichlid *Rhamphochromis esox* (Boulenger, 1908) was used for PCR positive controls (six per PCR plate; N = 114), whilst sterile molecular grade water (Fisher Scientific) substituted template DNA for No Template Controls, NTCs (six per PCR plate; N = 114). In the second PCR, Molecular Identification (MID) tags (unique 8-nucleotide sequences) and Illumina MiSeq adapter sequences were added to the amplified product. Two independent libraries were constructed, each containing 266 eDNA samples, 57 NTCs, and 57 positive controls. Sequencing was performed on an Illumina MiSeq using 2 ×300 bp V3 chemistry at Fera. The first sequencing run revealed human contamination across samples and in some PCR controls, therefore reactions prepared for the second sequencing run were sealed with mineral oil to minimise PCR contamination. Full details of the eDNA metabarcoding workflow are provided in Supporting Information: Appendix 3.

### BIOINFORMATIC PROCESSING

Illumina data was converted from raw sequences to taxonomic assignment using a custom pipeline for reproducible analysis of metabarcoding data: metaBEAT (metaBarcoding and eDNA Analysis Tool) v0.8 (https://github.com/HullUni-bioinformatics/metaBEAT). Bioinformatic data processing/analysis largely followed the workflow outlined by Hänfling *et al.* (2016), with minor modifications (see Supporting Information: Appendix 3 for details). To ensure reproducibility of analyses, the workflow has been deposited in the GitHub repository.

### DATA ANALYSIS

All downstream analyses were performed in the statistical programming environment R v.3.3.2. (R Core Team 2016). Data and R scripts have been deposited in the GitHub repository. Manipulation of the dataset produced by metaBEAT is described in Supporting Information: Appendix 4.

#### Detection thresholds and contamination

At present, there are no standard guidelines for eDNA analysis to indicate minimum number of positive eDNA samples or replicates required to class sites as species positive (Goldberg *et al.* 2016). Samples analysed by qPCR in this study were previously considered *T. cristatus* positive if one or more qPCR replicates gave a positive result (Biggs *et al.* 2015). We term this analysis qPCR NT (No Threshold). This inference of species presence is employed across many studies but may not be reliable or reproducible (Goldberg *et al.* 2016). More stringent qPCR thresholds reduced detection sensitivity for palmate newt *Lissotriton vulgaris* (Razoumowsky, 1789) (Smart *et al.* 2016) but may be necessary to ensure consistency and prevent false positives (Rees *et al.* 2014b). To facilitate comparison with current qPCR scoring (our NT interpretation) and eDNA metabarcoding, we applied a stringent qPCR threshold of >= 4/12 positive qPCR replicates to infer species presence, and termed the new analysis qPCR TA (Threshold Applied).

The raw eDNA metabarcoding dataset with no detection thresholds applied was termed metabarcoding NT (No Threshold). A second dataset was constructed to reduce the potential for false positives by application of a species-specific threshold: a species was only classed as present at a given site if its sequence frequency exceeded a species-specific threshold. Thresholds for each species were defined by analysing sequence data from PCR positive controls (N = 114) and identifying the maximum sequence frequency for a given species across all PCR positive controls (Supporting Information: Table S2). For example, the species-specific false positive sequence threshold for *T. cristatus* was 0.028% to omit all false detections in the PCR positive controls. The resultant dataset was termed metabarcoding TA (Threshold Applied).

We tested whether mineral oil reduced contamination by analysing the distribution of positive control sequences (*R. esox*) and human DNA in eDNA samples, and any DNA in NTCs, across both sequencing runs using binomial generalised linear mixed effects models (GLMMs) within the R package ‘lme4’ v1.1-12 (Bates *et al.* 2015). The response variable was presence-absence of contamination and explanatory variables were PCR plate (random effect) and sequencing run i.e. mineral oil sealed versus non-sealed (fixed effect). Human DNA may be present in eDNA samples as a real environmental signal or contaminant prior to PCR and thus may not be a true PCR contaminant. Consequently, contamination in eDNA samples was examined using several model permutations, where contamination comprised both cichlid and human DNA, cichlid DNA alone and human DNA alone. An information-theoretic approach using Akaike’s Information Criteria (AIC) to evaluate model fit was employed, where low AIC models are more parsimonious than high AIC models (Akaike 1973). Significance of the fixed effect in the model was tested by a Likelihood Ratio Test (LRT).

#### Comparison of eDNA methods for T. cristatus detection

We next tested the null hypothesis of no significant difference in sensitivity of qPCR and metabarcoding. Overall agreement between eDNA metabarcoding and qPCR for *T. cristatus* detection was measured using Cohen’s kappa coefficient (Cohen 1960), following which Pearson’s Chi-squared Test for Independence was used to test equality of *T. cristatus* detection between eDNA approaches.

Previously, Biggs *et al.* (2015) found qPCR score was an inconsistent predictor of *T. cristatus* abundance, where ponds with low scores had low newt counts but high scores did not correspond to large populations. qPCR score may only be proxy for the amount of DNA present rather than the number of individuals. The relationship between read count and qPCR score has not been examined previously, and whether read production is indicative of DNA concentration remains unknown. We hypothesised samples with higher qPCR score would have increased *T. cristatus* read count. First, the average number of *T. cristatus* reads produced by eDNA metabarcoding per qPCR score (1-12 out of 12) was calculated. A Spearman Rank Correlation was then used to test for a relationship between average read count and qPCR score.

Following data exploration (see Supporting Information: Appendix 4), a negative binomial GLMM was used to counter overdispersion and improve model fit. The GLMM examined read count in relation to qPCR score, accounting for other variables that may affect metabarcoding signal strength. Variation in *T. cristatus* read count was examined using the proportion of *T. cristatus* reads within the total number of reads produced for each eDNA sample as the response variable. Sequencing run and PCR plate were considered random effects and all other explanatory variables as fixed effects (qPCR score, sample degradation, sample inhibition, post-PCR eDNA concentration). Presence-absence of sample degradation and inhibition was determined by qPCR in 2015 using methodology outlined by Biggs *et al.* (2015). Model fit was again evaluated using AIC and significance of fixed effects in the model was tested with stepwise backward deletion of terms from the model informed by LRTs. All values were bound in a new data frame and model results plotted for evaluation using the R package ‘ggplot2’ v 2.1.0 (Wickham, 2009).

#### Cost and investigator effort

Cost of materials, and investigator effort and salary (assuming hourly rate of £21.20) were calculated for eDNA samples but estimates do not include travel to sampling sites, procedural controls, qPCR standards, or consumables and reagents required for assay optimisation. Time required to perform PCR for metabarcoding and qPCR was estimated assuming available machinery to run four PCR plates in parallel and one qPCR plate.

## Results

### TARGETED qPCR AND EGG SEARCHES

Targeted qPCR detected *T. cristatus* in 253 (49.80%) samples analysed by Fera (N = 508). Of 255 (50.20%) samples that were negative, one was inhibited and nine were degraded. qPCR and egg searches produced consistent results for 297 (58.47%) ponds, with 51 (10.04%) positive and 246 (48.43%) negative ponds by both methods. Of the 211 ponds where there was disagreement between methods, 202 (39.76%) were qPCR positive but negative by egg searches, and 7 (1.38%) were positive with egg searches but qPCR negative. Of 24 samples analysed by ADAS, 12 (50.00%) were qPCR negative and 12 (50.00%) were qPCR positive for *T. cristatus*. No egg search data were available for these ponds.

### VERTEBRATE METABARCODING

The *in silico* and *in vitro* primer validation confirmed that *T. cristatus*, and other native UK amphibians tested, can be reliably amplified and identified with the chosen assay (Supporting Information: Appendix 5, Fig. S1). Furthermore, the *in silico* approach showed that the majority of all UK vertebrates can be amplified (see Appendix 5 for details). Both sequencing runs had comparable yield and sequencing quality score; summary statistics for each sequencing run and read counts for taxonomic assignment levels are provided in Supporting Information: Appendix 5 (Tables S3, S4). A full summary of sequence read count data is also given in Supporting Information: Appendix 5 (Table S5). eDNA metabarcoding identified a combined total of 60 species (Supporting Information: Appendix 5, Fig. S2) across both sequencing libraries, with 375,954 and 508,879 sequences assigned to *T. cristatus* from each library. Analyses of overall pond species compositions inferred by eDNA metabarcoding (Supporting Information: Appendix 5, Fig. S3, Table S6) are reported separately (Harper *et al.* in preparation).

All samples (N=532) were sequenced and of 57 samples that did not produce visible PCR bands, nine generated sequence reads. Notably, the 57 samples were not inhibited or degraded at time of qPCR. Weak PCR bands were observed in some NTCs therefore all PCR controls were sequenced (Supporting Information: Appendix 5, Fig. S4). Six NTCs contained *T. cristatus* DNA but only one exceeded 100 *T. cristatus* reads (307/330 reads). Twelve other sources occurred in NTCs (Supporting Information: Appendix 5, Table S7); seven occurred in more than one NTC and eight had high maximum read counts (> 100 reads).

Contamination of NTCs (any DNA) and environmental samples (cichlid/human DNA) was observed (Supporting Information: Appendix 5, Figs. S4, S5, S6). Read counts of NTC contaminants were reduced between sequencing runs with the addition of mineral oil to PCR reactions included on the second sequencing run (Supporting Information: Appendix 5, Figs. S4, S5, S6) but this reduction was not statistically significant (GLMM: *Χ*^2^1, *F*1 = 1.941, *P* > 0.05). Mineral oil did not reduce human DNA signal in environmental samples between sequencing runs either (GLMM: *Χ*^2^1, *F* _1_ = 3.591, *P* > 0.05); however, it did reduce human DNA in combination with cichlid DNA (GLMM: *Χ*^2^1, *F*_1_ = 21.143, *P* < 0.01), and cichlid DNA contamination alone (GLMM: *Χ*^2^1, *F*_1_ = 6.978, *P* < 0.05) of environmental samples.

### eDNA METABARCODING VS qPCR FOR *T. CRISTATUS* DETECTION

*T. cristatus* detection by metabarcoding NT (34.21%) was less sensitive than qPCR NT (49.81%) but marginally higher than qPCR TA (32.71%) (N = 532 ponds, Fig. 2). Metabarcoding TA had lower detection efficiency (28.01%) and failed to detect *T. cristatus* in 116 and 25 ponds where the species was detected by qPCR NT and qPCR TA respectively. Nonetheless, both molecular approaches attained higher *T. cristatus* detection than daytime egg searches (11.46%) in 506 ponds where all three approaches were implemented.

**Figure 2.**
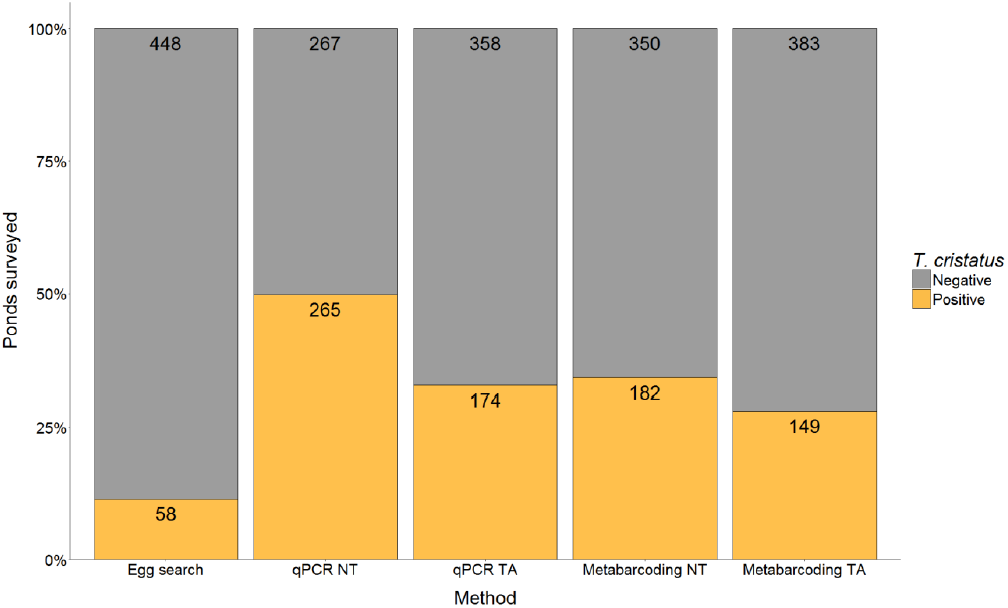
Comparison of survey methodology for *T. cristatus* detection in freshwater ponds across the UK. Bars represent proportion of positive and negative *T. cristatus* ponds by each method with frequency displayed on bars.

Overlap between survey methods for positive *T. cristatus* ponds (N = 277), and unique detections by each method is summarised in Fig. 3. Negative *T. cristatus* ponds (N = 229) are examined in combination with species positive ponds in Supporting Information: Appendix 5 (Table S8). Each survey method detected the species in ponds where other methods failed. Despite lower *T. cristatus* detection efficiency, egg searches detected the species in six ponds where it went undetected by qPCR and metabarcoding. Metabarcoding NT and metabarcoding TA revealed *T. cristatus* in seven ponds which other methods did not, whilst qPCR NT and qPCR TA detected *T. cristatus* in 33 ponds unique to other methods. All methods detected *T. cristatus* in 32 ponds, and both metabarcoding and qPCR identified *T. cristatus* in 86 ponds. Disagreement between molecular methods was more likely when samples were positive rather than negative by qPCR. Without thresholds, 39.25% of qPCR positive ponds (N = 265) were negative by metabarcoding, but 7.87% of qPCR negative ponds (N = 267) were positive by metabarcoding. With thresholds, 29.31% of qPCR positive ponds (N = 174) were negative by metabarcoding, whereas 7.26% of qPCR negative ponds (N = 358) were positive by metabarcoding.

**Figure 3.**
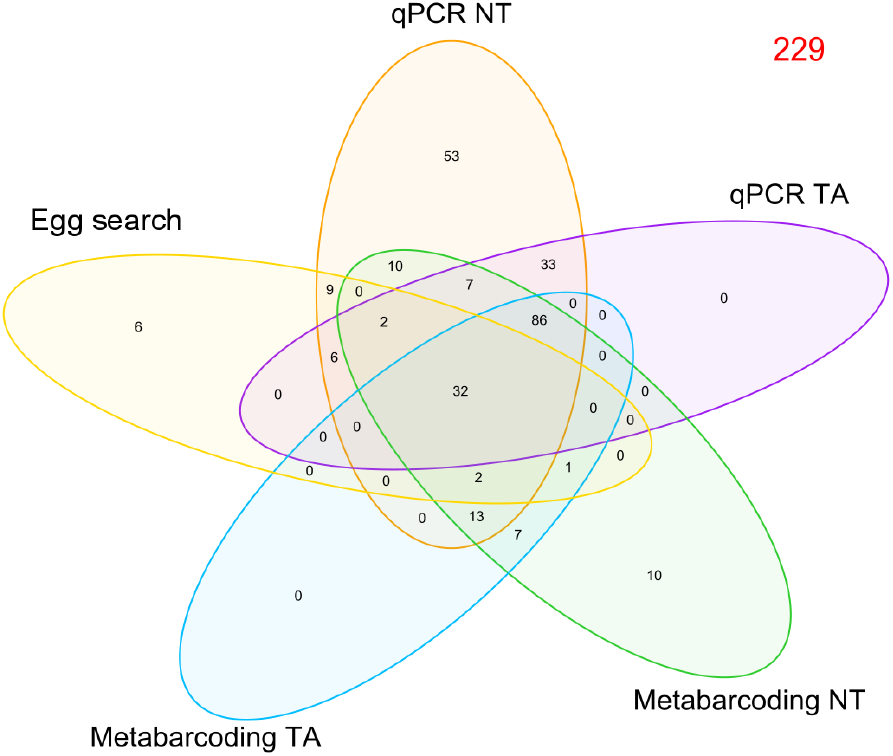
Venn diagram which summarises the number of positive *T. cristatus* detections (N = 277) by each method (egg search, qPCR NT, qPCR TA, metabarcoding NT, and metabarcoding TA), and overlap in *T. cristatus* detection 797 between methods for 506 ponds where all methods were applied. Negative *T. cristatus* detections (N = 229) are 798 highlighted in red.

Agreement between eDNA approaches is summarised in Table 1. Agreement was strongest between eDNA approaches when the qPCR detection threshold was applied, irrespective of whether the metabarcoding detection threshold was applied. Metabarcoding (NT or TA) and qPCR TA did not significantly differ in their detection of *T. cristatus* (Table 1). An identical positive correlation was observed between qPCR score and the average number of *T. cristatus* reads obtained for samples belonging to each qPCR score (*r*_*s*_ = 0.648, df = 11, *P* < 0.05), regardless of threshold application to the metabarcoding data. Despite some inconsistency across qPCR scores, samples with a higher qPCR score generally had more *T. cristatus* reads, supportive of a relationship between metabarcoding and abundance of eDNA from single species. Notably, metabarcoding produced *T. cristatus* reads for qPCR NT and qPCR TA negative samples, but the *T. cristatus* metabarcoding signal of these (qPCR NT_negative_ = 2639 reads max., qPCR TA_negative_ = 3075 reads max.) was much lower than samples with higher qPCR score (max. 65,325 reads; Supporting Information: Appendix 5). Further examination of the relationship between qPCR score and metabarcoding TA revealed qPCR score and post-PCR eDNA concentration of samples also influenced the proportion of *T. cristatus* reads i.e. relative *T. cristatus* sequence read production (Table 2). A significant positive relationship was observed between qPCR score and the proportion of *T. cristatus* reads within total reads per sample (*P* < 0.001) (Fig. 4a). Conversely, post-PCR eDNA concentration had a significant negative influence on the proportion of *T. cristatus* reads (*P* < 0.001), where read proportion decreased as post-PCR eDNA concentration increased (Fig. 4b).

**Table 1:**
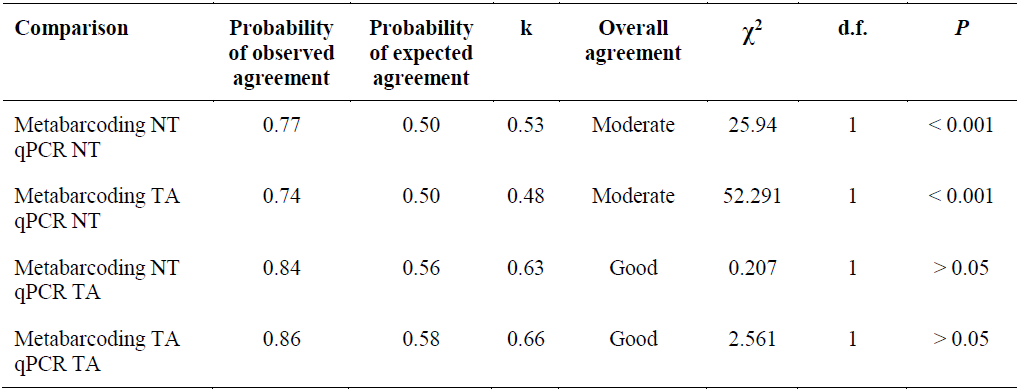
Summary of analyses testing for agreement between eDNA approaches, with threshold applied (TA) and no threshold (NT), for *T. cristatus* detection. Cohen’s kappa coefficient (k) represents strength of agreement between methods (1 = 100%). Pearson’s Chi-squared Test for Independence tested whether methods significantly differed for *T. cristatus* detection.

**Table 2:**
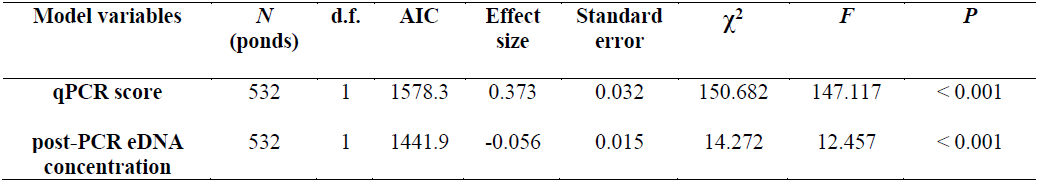
Summary of analyses testing for variation in proportion of *T. cristatus* sequence reads in a sample produced by eDNA metabarcoding, attributable to qPCR score or post-PCR eDNA concentration. Test statistic is for LRT used.

| Model variables | *N*<br>(ponds) | d.f. | AIC | Effect<br>size | Standard<br>error | $\chi^2$ | *F* | *P* |
| --- | --- | --- | --- | --- | --- | --- | --- | --- |
| qPCR score | 532 | 1 | 1578.3 | 0.373 | 0.032 | 150.682 | 147.117 | < 0.001 |
| post-PCR eDNA<br>concentration | 532 | 1 | 1441.9 | -0.056 | 0.015 | 14.272 | 12.457 | < 0.001 |

**Figure 4.**
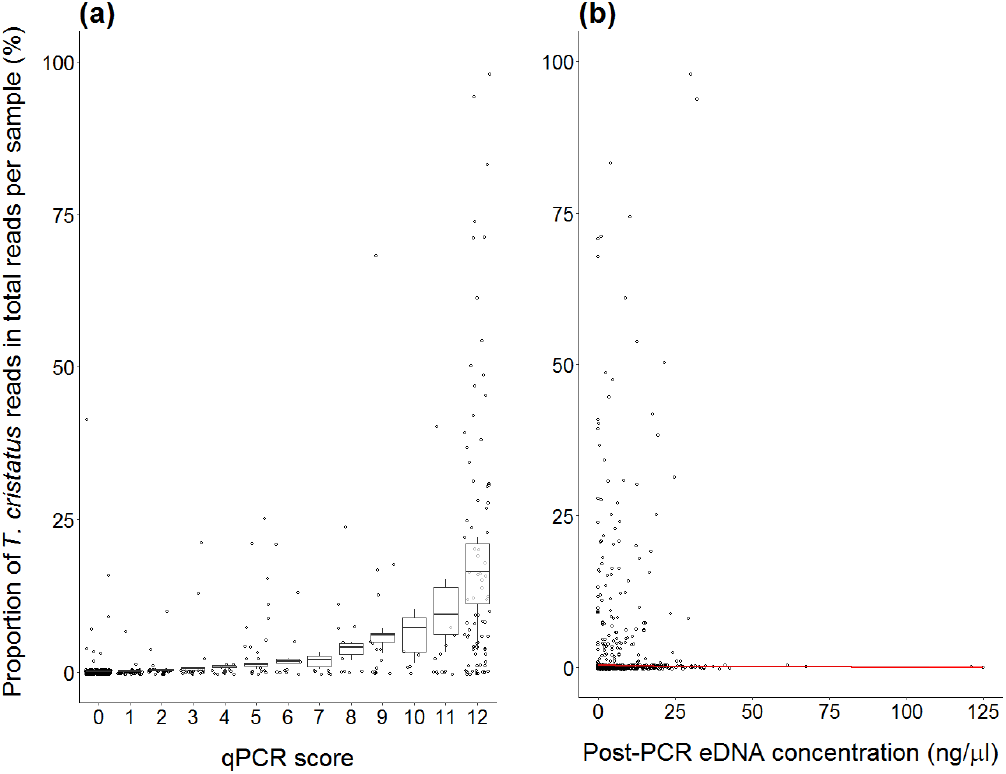
Relationship between fixed effects (qPCR score, post-PCR eDNA concentration) and response variable (proportion of *T. cristatus* reads) in eDNA samples, as predicted by the negative binomial GLMM. The 95% CIs, as 802 calculated using the predicted proportions, and standard error for these predictions are given for each relationship. 803 The observed data (points) are also displayed against the predicted relationships (boxes, line). The proportion of *T. cristatus* reads within eDNA samples increased as qPCR score increased **(a)**. Conversely, the proportion of *T. cristatus* reads decreased as post-PCR eDNA concentration increased **(b)**.

### COMPARISON OF METHOD COST AND INVESTIGATOR EFFORT

Cost and investigator effort for both eDNA approaches were comparable. Metabarcoding was marginally more expensive (£3 per pond) than qPCR, but used 1 day less of investigator effort. A full breakdown of expenditure per pond is given in Supporting Information: Appendix 5 (Table S9) and summarised in Fig. 5.

**Figure 5.**
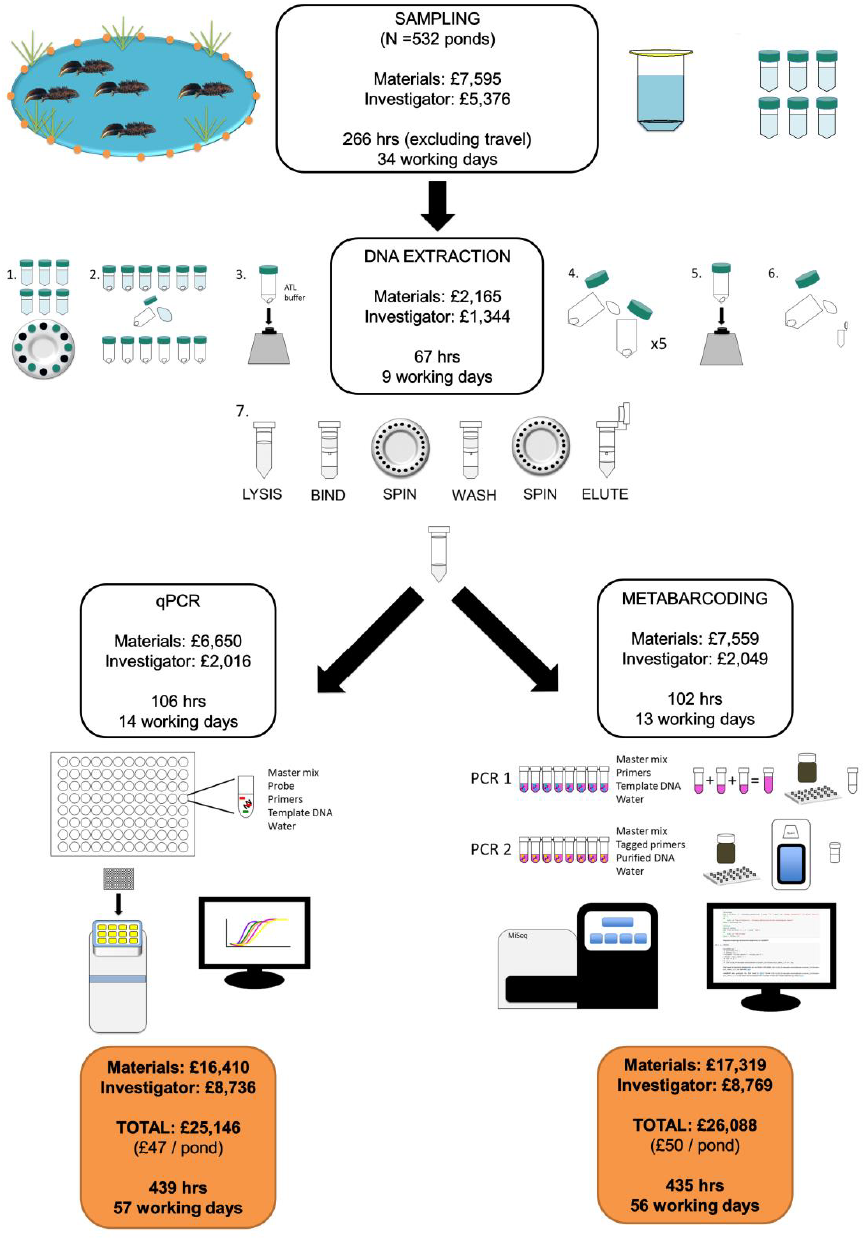
Cost and investigator effort required for targeted qPCR of *T. cristatus* and eDNA metabarcoding of vertebrate communities from pond water samples.

## Discussion

We have demonstrated eDNA metabarcoding is a highly sensitive tool for monitoring *T. cristatus* alongside the wider biological community, corroborating other comparisons of eDNA metabarcoding and qPCR for single-species monitoring (Lacoursière-Roussel *et al.* 2016; Schneider *et al.* 2016). Despite reduction in single-species detection, eDNA metabarcoding revealed a wealth of biodiversity information and could enable more effective freshwater monitoring networks and better understanding of community structure and ecosystem function alongside *T. cristatus* monitoring (Biggs, von Fumetti & Kelly-Quinn 2016). However, both eDNA approaches have advantages and drawbacks which must be considered for design and implementation of biodiversity monitoring programmes.

### SINGLE-SPECIES DETECTION BY qPCR AND METABARCODING

A direct comparison of sensitivity between qPCR and metabarcoding is not straightforward: stochasticity in qPCR largely occurs during amplification (volume of template DNA and technical replication), whereas stochastic variation during metabarcoding arises through PCR amplification and sequencing (depth and replication) (Kelly *et al.* 2014; Thomsen *et al.* 2016; Deiner *et al.* 2017). In our study, 12 independent qPCR replicates were performed for each sample but due to limited resources, metabarcoding was based on three pooled PCR replicates which were sequenced once only. Therefore, to enable a fair comparison between methods in terms of PCR effort, a threshold of >= 4/12 positive replicates (qPCR TA) was applied to the qPCR data. Detection sensitivity was most similar between methods with the qPCR threshold and without the metabarcoding threshold. Both eDNA metabarcoding and qPCR displayed reduced *T. cristatus* detection when thresholds were applied; although, this may reflect reduced false positive detections rather than decreased sensitivity. Lower sensitivity of the eDNA metabarcoding approach used here may also stem from sample degradation during long-term storage. The samples used were stored for more than 12 months at -80°C before metabarcoding. However, long-term storage and continual freeze-thawing of samples may allow aggregation of inhibitory substances which impair PCR amplification and cause false negatives (Takahara, Minamoto & Doi 2015).

Despite lower sensitivity, strength of eDNA signal produced by metabarcoding was correlated with that of qPCR, where both *T. cristatus* average read count and read proportion broadly increased with qPCR score of eDNA samples. The correlation was inconsistent though, where high average or proportional *T. cristatus* read count did not always correspond to high qPCR score. Biggs *et al.* (2015) also found a variable positive association between qPCR and *T. cristatus* counts, where high qPCR score did not always correlate with high counts. Quantitative data on eDNA concentration are needed to examine the performance of each eDNA approach in relation to the amount of eDNA present, and whether these tools can reliably estimate species abundance. This data can be obtained with highly sensitive qPCR assays, and inclusion of internal DNA standards in sequencing runs for metabarcoding (Ushio *et al.* 2017). Nonetheless, our results suggest performance of metabarcoding and qPCR are linked and influenced by external factors. Evans *et al.* (2016) suggested the relative abundance and biomass of a species interact to exert a combined effect on eDNA production rate and subsequent metabarcoding detection. The abundance, biomass, and distribution of *T. cristatus* (Biggs *et al.* 2015), as well as shedding rate, environmental factors, and eDNA transport (Goldberg *et al.* 2016; Buxton *et al.* 2017), may all influence detection and concentration of eDNA, and inferences made using qPCR and metabarcoding.

The comparison between qPCR and metabarcoding must also be examined in context of the sequencing effort. Here, we sequenced a large number of samples (380 including PCR controls) per run to provide a realistic cost scenario for routine monitoring. Yet, metabarcoding sensitivity would likely improve with an increase in read depth per sample (Kelly *et al.* 2014). In order to directly compare eDNA signal production by these approaches, it may be necessary to perform sequencing replicates to verify true positives where rare species are expected and generate an “eDNA metabarcoding score” system similar to qPCR (Brandon-Mong *et al.* 2015; Port *et al.* 2016; Civade *et al.* 2016; Thomsen *et al.* 2016). PCR and sequencing replication in metabarcoding may enhance species detection probability through improved amplification of low abundance or highly degraded DNA (Ficetola *et al.* 2015; Port *et al.* 2016) that is readily amplified by qPCR (Lacoursière-Roussel *et al.* 2016).

Similarly, sequencing of independent biological replicates, opposed to pseudoreplicates from a single water sample, may improve detection and minimise false negatives produced by eDNA metabarcoding (Andruszkiewicz *et al.* 2017; Bálint *et al.* 2017). Currently, 90 mL (6 × 15 mL sampled from 600 mL) water is sampled during *T. cristatus* eDNA survey, followed by ethanol precipitation (Biggs *et al.* 2015). While this may be appropriate for highly-sensitive targeted qPCR, larger water volumes and filtration may be required to capture eDNA from less abundant vertebrates and characterise community diversity (Shaw *et al.* 2016b). Additionally, eDNA from different species, and individuals within species, can be unevenly distributed throughout water bodies and may be concentrated in particular areas (Biggs *et al.* 2015; Hänfling *et al.* 2016; Evans *et al.* 2017a), thus sampling strategies must be carefully designed to ensure eDNA samples are representative of biodiversity present.

Metabarcoding assays are also susceptible to problems from taxon bias, DNA swamping and bioinformatics related problems (Taberlet *et al.* 2012; Kelly *et al.* 2014; Shaw *et al.* 2016b). Potential reduction in sensitivity of passive community sequencing versus targeted qPCR may relate to the performance of metabarcoding primers for target species. During metabarcoding, DNA from rare species may be masked by highly abundant species (Schneider *et al.* 2016), or under-represented due to disproportionate eDNA shedding rates across species and preferential amplification of other species (Kelly *et al.* 2014). PCR-free workflows (i.e. shotgun sequencing) eliminate this bias through indiscriminate sequencing; however, this is unsuitable for conservation projects with target species as a mass of uninformative data are produced, and too costly for routine monitoring schemes (Shaw *et al.* 2016b; Valentini *et al.* 2016). We found *T. cristatus* read proportion was negatively associated with post-PCR concentration of eDNA samples. As a positive relationship was not observed, this would suggest PCR amplification with our selected marker and primers was not biased toward our focal species. However, we cannot conclude that our metabarcoding assay was free of primer bias as post-PCR concentration of eDNA samples can be influenced by PCR stochasticity.

Multiple markers (e.g. COI, CytB, 12S, 16S) are increasingly used in eDNA metabarcoding to cast a wider net of species detection and minimise primer bias (Evans *et al.* 2016, 2017a; Valentini *et al.* 2016; Hänfling *et al.* 2016; Shaw *et al.* 2016a). Using markers from both mitochondrial and nuclear genes may reduce bias associated with specific genes or primers, and provide greater taxonomic resolution (Kelly *et al.* 2014). Furthermore, multiple markers of different lengths may enhance understanding of eDNA persistence and state, and species location. Long barcodes bind to stable DNA that has been recently deposited by species (Hänfling *et al.* 2016), and may reduce false negatives whilst increasing taxonomic resolution and accuracy (Kelly *et al.* 2014; Valentini *et al.* 2016; Shaw *et al.* 2016a). In contrast, short barcodes (such as 12S used here) challenge sequencers and bioinformatics tools (Taberlet *et al.* 2012; Shaw *et al.* 2016a), but readily amplify short, degraded DNA fragments that persist longer and possibly disperse further in water bodies, improving probability of detection (Hänfling *et al.* 2016). It is possible that metabarcoding detection rates could be improved by using group-specific metabarcoding primers for amphibians, such as the ‘batra’ set recently designed by Valentini *et al*. (2016). More specific primers could increase relative coverage of *T. cristatus,* providing more comparable detection rates to qPCR. This is worth investigating, but with the caveat that group-specific primers obviously restrict the biodiversity information that can be gained from an ecosystem.

### FALSE NEGATIVES

This study did not aim to evaluate sensitivity of molecular methods against standard *T. cristatus* survey methodologies. Egg searches were used to detect false negatives produced by qPCR and metabarcoding and in doing so, revealed some interesting results. Biggs *et al.* (2015) previously demonstrated qPCR had higher detection rate than egg searches (as well as torchlight, netting, and bottle trapping), but here we show this also holds true for metabarcoding. Importantly, absence of eggs does not infer absence of adults, and this method is highly dependent on weather conditions and water clarity (Rees *et al.* 2014a; Biggs *et al.* 2015). Despite considerably higher detection rate of both eDNA approaches, eggs were recorded in a small number of ponds that were eDNA negative. eDNA analysis can incorrectly infer absence or low abundance of species if inhibition or interference from non-target DNA has occurred (Goldberg *et al.* 2016). Alternatively, eDNA false negatives may have been a by-product of sampling strategy and effort for *T. cristatus*. Larger water volumes and/or more biological replication instead of pseudoreplication (established *T. cristatus* eDNA sampling strategy) may improve detection (Lopes *et al.* 2016; Andruszkiewicz *et al.* 2017; Bálint *et al.* 2017). All methods revealed *T. cristatus* in ponds where other approaches failed, emphasising that these species monitoring tools are complementary and should be used in combination to achieve maximum detection probability. However, integrative strategies combining molecular and conventional tools are often not cost-efficient for most applications.

### FALSE POSITIVES

False positives may arise from field contamination and eDNA transport in the environment particularly by waterfowl (Shaw *et al.* 2016a). eDNA is retained by predators, discarded in faeces, and transported by anthropogenic activity, combined with natural water currents and flow (Hänfling *et al.* 2016). In the laboratory, PCR-accumulated and sequencing error, including primer mismatch (Andersen *et al.* 2012) and ‘tag jumps’ (Schnell, Bohmann & Gilbert 2015), can induce misassignment leading to false positives, cross-contamination between samples, or laboratory contamination (Andruszkiewicz *et al.* 2017).

False positives can be modelled and estimated using Site Occupancy Modelling of metabarcoding data (Ficetola *et al.* 2015) or risk of false positives minimised using a sequencing threshold i.e. the number of sequence reads required for a sample to be species positive (Hänfling *et al.* 2016; Civade *et al.* 2016; Evans *et al.* 2017a). However, such thresholds can reduce detection of rare species, a primary goal of this study, and may fail where false and true positives occur at similar frequency (Hänfling *et al.* 2016). Instead, we calculated species-specific sequence thresholds to more accurately control for false positives in our dataset without compromising *T. cristatus* detection.

In our study, human DNA occurred at high frequency and abundance; this may have been a true environmental signal from pond water, or real contaminant as encountered in other metabarcoding research (Port *et al.* 2016; Valentini *et al.* 2016; Thomsen *et al.* 2016). Blocking primers can prevent amplification of abundant non-target DNA like human (Valentini *et al.* 2016) but may fail (Thomsen *et al.* 2016) or prevent amplification of target taxa (Port *et al.* 2016). Alongside human, other aquatic and terrestrial vertebrate DNA occurred at high frequency in NTCs, although these were not removed by addition of mineral oil. An even stricter forensic laboratory set-up, such as that employed for ancient DNA (aDNA), should be adopted to ensure data robustness. Positive and negative controls should be included at each stage of metabarcoding workflows to monitor contamination (Deiner *et al.* 2017). However, preventive measures inevitably increase research cost and some degree of contamination is unavoidable in metabarcoding (Kelly *et al.* 2014; Brandon-Mong *et al.* 2015; Port *et al.* 2016; Thomsen *et al.* 2016).

Our results also highlight the importance and impact of qPCR thresholds when inferring species presence-absence. Similar to Smart *et al.* (2016), we found a stringent qPCR threshold reduced detection sensitivity. As yet, no guidance exists to indicate how many samples or replicates must be positive to class a site as species-positive (Goldberg *et al.* 2016; Smart *et al.* 2016) but clearly this must be addressed to improve standardisation and reproducibility of eDNA research. Importantly, less stringent thresholds (and false positives inherent to these) are somewhat precautionary and may better protect *T. cristatus* by preventing development. Therefore, whilst reduction or removal of false positives is desirable, detection thresholds must not compromise protection of threatened species either. Until a suitable threshold can be established, it may be more appropriate to re-analyse samples which yield one positive qPCR replicate to prevent false positives (Rees *et al.* 2014b; Goldberg *et al.* 2016).

### COST AND INVESTIGATOR EFFORT

Cost efficiency combined with the overarching aim of a monitoring or conservation programme should always be considered. We found eDNA metabarcoding was more costly than qPCR but both approaches required similar investigator effort. qPCR scales to the number of samples being processed (Schneider *et al.* 2016) whereas metabarcoding has fixed costs including reagent kit for HTS platform (Bálint *et al.* 2017). eDNA metabarcoding becomes more cost-efficient as more samples are processed (Bálint *et al.* 2017) but fewer replicates would reduce qPCR cost (Davy *et al.* 2015; Smart *et al.* 2016). Cost of eDNA monitoring is influenced by sample size, methods, replication, laboratory, statistical power and occupancy modelling (Davy *et al.* 2015; Evans *et al.* 2017b). Consequently, cost is proportional to project requirements (Davy *et al.* 2015) and will vary depending on choice of qPCR or metabarcoding workflow. Whilst qPCR is established technology that has reached its price ceiling, HTS is relatively new technology and prices will continue to drop, meaning higher sample throughput and more technical replication will be possible. We therefore argue that metabarcoding will become more cost-efficient in the long-term, providing more data at lower cost and comparable sensitivity to qPCR. However, where samples cannot be processed in large batches, qPCR may retain cost-efficiency.

## Conclusion

eDNA metabarcoding holds promise for holistic biodiversity monitoring of freshwater ponds as opposed to targeted qPCR for flagship or indicator species such as *T. cristatus*. Metabarcoding can reveal entire species assemblages from environmental samples without prior ecosystem information, and provide broad-scale distribution data for multiple species simultaneously.Nonetheless, the method at present appears to be less sensitive than qPCR for single-species monitoring, and species detection by molecular and conventional methods was incongruent. Comprehensive study of the influence of water volume, eDNA capture and extraction method, and sample storage on single-species and community detection in lentic and lotic systems is required. Minimising the risk of false positives and contamination remains a pressing issue in metabarcoding, and standard contamination measures (Goldberg *et al.* 2016) may be insufficient for analysis of vertebrate assemblages. Currently, cost and investigator effort required for metabarcoding and qPCR are broadly equivalent, but reduced sequencing costs may level the playing field. We conclude that eDNA metabarcoding is not yet a replacement for targeted qPCR and conventional survey but rather another tool in the ecologist toolbox. Ultimately, choice of monitoring tool(s) is specific to aims of each conservation project. At present, qPCR retains sensitivity for *T. cristatus* populations of all sizes, regardless of sample number processed. Under a realistic conservation monitoring scenario, where funding is limited and samples must be processed in large batches, metabarcoding may suffer from false negatives due to reduced sequencing depth and replication. However, in many cases, the biodiversity information generated by this approach, and its implications for community ecology and conservation, will eclipse lower sensitivity. This passive screening approach would be most effective for initial survey of water bodies to generate broad-scale multi-species distribution data. This holistic data can then inform best use of funding and time for targeted species-specific survey.

## Acknowledgements

This work was funded by University of Hull. We would like to thank Jennifer Hodgetts (Fera) for assisting with sample collection and data dissemination, and Jianlong Li (University of Hull) for primer design and advice on laboratory protocols. Furthermore, Barbara Mabel, Elizabeth Kilbride (University of Glasgow), Andrew Buxton and Richard Griffiths (DICE, University of Kent) provided tissue samples for primer validation and Sanger sequencing to supplement the reference database.

## Author contributions

B.H., L.R.H., L.L.H and N.B. conceived and designed the study. H.C.R., K.C.G., and N.B. contributed samples for processing. H.C.R. performed extractions under licence from Natural England and qPCR for eDNA samples from private contracts at The University of Nottingham.N.B. performed qPCR for eDNA samples from Natural England’s Great Crested Newt Evidence Enhancement Program. L.R.H. performed remaining laboratory work and analysed the data. I.P.A. and E.L. offered advice on and supervised sequencing. C.H. assisted with bioinformatics analysis.P.B. and S.P. contributed datasets for analysis. L.R.H. wrote the manuscript, which all authors revised.

## Data accessibility

Raw sequence reads have been archived on the NCBI Sequence Read Archive (Bioproject: PRJNA417951; SRA accessions: SRR6285413 - SRR6285678). Jupyter notebooks, R scripts and corresponding data are deposited in a dedicated GitHub repository (https://github.com/HullUni-bioinformatics/Harper_et_al_2018”bioinformatics/Harper_et_al_2018) which has been permanently archived (https://doi.org/10.5281/zenodo.1188709).

